# Multiple-biological matrices metabolomics identified new metabolite biomarkers for the precise diagnosis of pancreatic cancer and associated tissue metastasis

**DOI:** 10.1101/2020.03.09.983239

**Authors:** Xialin Luo, Jingjing Liu, Huaizhi Wang, Haitao Lu

**Author notes:** Corresponding authors Haitao Lu., Ph.D., Professor, Shanghai Jiao Tong University, Huaizhi Wang M.D., Ph.D., Professor., University of Chinese Academy of Sciences.

## Abstract

**Purpose:** To improve clinical diagnosis and enhance therapeutic outcome, we figure out to identify and validate metabolite biomarkers from the plasma samples of patients with pancreatic cancer that can easily, sensitively and efficiently diagnose the onsite progression, and metastasis of the disease.

**Experimental Design:** We employed the newly developed precision-targeted metabolomics method to validate that many differential metabolites have the capacity to markedly distinguish patients with pancreatic cancer from healthy controls. To further enhance the specificity and selectivity of metabolite biomarkers, a dozen tumor tissues from PC patients and paired normal tissues were used to clinically validate the biomarker performance.

**Results:** We eventually verified five new metabolite biomarkers in plasma (creatine, inosine, beta-sitosterol, sphinganine and glycocholic acid), which can be used to readily diagnose pancreatic cancer in a clinical setting. Excitingly, we proposed a panel biomarker by integrating these five individual metabolites into one pattern, demonstrating much higher accuracy and specificity to precisely diagnose pancreatic cancer than conventional biomarkers (CA125, CA19-9, CA242 and CEA); Moreover, we characterized succinic acid and gluconic acid as having a great capability to monitor the progression and metastasis of pancreatic cancer at different stages.

**Conclusions:** Taken together, this metabolomics method was used to identify and validate metabolite biomarkers that can precisely and sensitively diagnose the onsite progression and metastasis of pancreatic cancer in a clinical setting. Furthermore, such effort should leave clinicians with the correct time frame to facilitate early and efficiently therapeutic interventions, which could largely improve the five-year survival rate of PC patients.

## Introduction

Pancreatic cancer (PC) is a malignant tumour in the clinic; although the disease occurrence rate is quite low, it is still likely to become the second leading cause of cancer-related death worldwide due to the extremely low 5-year survival rate of less than 5% ^1-3^. Basically, the high mortality of patients with PC is attributed to the complex pathogenesis, and latency leads to a severe lack of early diagnostic methods. Currently, surgical resection is the main treatment measure to intervene in PC, but over 80% of PC patients already have suffered multiple organ metastases when the disease is clinically diagnosed; unfortunately, these patients have lost the opportunity to cure the disease^2^. In the clinic, serum carbohydrate antigen 19-9 (CA19-9), CA125, CA242 and carcinoembryonic antigen (CEA) are regarded as common tumour markers to phenotype PC^4^, but they are not highly recommended as biomarkers to efficiently diagnose PC due to their low sensitivity and specificity ^5,6^. Although there is increasing data to show that mutations in KRAS, TP53 and SMAD4 can drive pancreatic tumourigenesis^7, 8^, PC pathogenesis is too complex to better understand the molecular mechanisms of onsite progression and metastasis. However, the discovery and identification of novel biomarkers that can sensitively and specifically diagnose the onsite progression and metastasis of PC in a clinical setting is urgently needed. In addition, mechanism-associated biomarkers are also needed in a timely manner to elucidate the pathogenesis of PC at the molecular level^9^, which will largely assist in the development of new strategies to timely diagnose the onsite progression and metastasis of PC, as well as to assess therapeutic outcomes by monitoring changes in biomarkers.

Cancers are typical metabolic diseases whose development indeed incurs substantial metabolic alterations^1,10^. The Warburg effect is defined as the overexpression of aerobic glycolysis in cancer cells that can shunt glycolytic intermediates into multiple metabolic pathways that generate nucleotides, lipids, and amino acids^11^. There is evidence to show that glutamine facilitates the progression of pancreatic cancer through a Kras-regulated metabolic pathway, and metabolomics has been used to preliminarily explore biomarkers to diagnose pancreatic cancer^13^. However, the limitations of metabolomics methods and sampling and clinical-wide metabolite biomarkers for the efficient and specific diagnosis of PC are still highly required in clinical practice. Here, we used a newly developed precision-targeted metabolomics method^14^ to analyse the plasma samples of patients with PC and healthy controls to precisely discover and characterize the most sensitive and specific diagnostic biomarkers and progressive biomarkers for monitoring the onsite progression and metastasis of PC. This new targeted metabolomics method has the capacity to profile 240 small molecule metabolites with important biological functions, with wide coverage of numerous important metabolic pathways involving the tricarboxylic acid cycle (TCA cycle), glycolysis, the pentose phosphate pathway (PPP), amino acid metabolism, etc., which have been reported to be closely associated with the pathogenesis of PC^15,16^. Our effort is first to identify biomarkers that can readily diagnose PC in a clinical setting. In addition, pathological latency often leads to metastasis of PC, and few biomarkers have been characterized for the early diagnosis of metastasis ^17^. Thus, our efforts also aim to identify biomarkers that enable the monitoring of progression and metastasis. Altogether, identifying diagnostic biomarkers can achieve the early diagnosis of PC as well as efficiently monitor the progression and metastasis of PC, by which clinicians could deliver therapeutic interventions at the right time, enabling a significant improvement in therapeutic outcomes by enhancing the 5-year survival rate. Moreover, mechanism-associated biomarkers might be captured to better understand the pathogenesis of PC from a metabolic perspective.

## Materials and methods

### Clinical samples

Plasma samples were collected from 60 patients diagnosed with pancreatic cancer and a cohort of 60 healthy volunteers from Southwest Hospital (Chongqing, China) from July 2017 to November 2018. The clinical data, including age, gender, and clinical pathological features, were all recorded in detail. The PC plasma samples were collected prior to surgical resection and stored at −80 °C for experimental use. Six paired pancreatic cancer tissue (PCT) and adjacent noncancerous tissue (ANT) samples were also provided by Southwest Hospital (Chongqing, China). This research was approved by the Southwest Hospital Institutional Ethics Review Board, and all participants provided written informed consent.

### Cell culture

The normal human pancreatic duct cell line hTERT-HPNE, the human primary pancreatic adenocarcinoma cell line BxPC-3 and the liver-metastatic pancreatic cell line SU.86.86 were purchased from the American Type Culture Collection (ATCC). The BxPC-3 and SU.86.86 cell lines were maintained in RPMI 1640 medium supplied with 10% foetal bovine serum and 1% penicillin-streptomycin. hTERT-HPNE was grown in one mixed medium that contained 25% medium M3 Base (Incell Corp. Cat. M300F-500), 75% glucose-free DMEM (Sigma Cat. D5030 with an additional 2 mM L-glutamine and 1.5 g/L sodium bicarbonate), 5% FBS, 10□ng/ml human recombinant EGF, 5.5□mM glucose, and 750 ng/ml puromycin. All cells were maintained at 37 °C in a humidified atmosphere containing 5% CO^2^.

### Sample preparation

Frozen plasma samples are thawed and homogenized, and then 100 µL of the supernatants are transferred into a 1.5 mL microcentrifuge tube, mixed with 400 µL of iced-cold acetonitrile and placed on ice to precipitate the proteins. Next, the mixture was centrifuged at 20,000 g for 10 min at 4 °C. To remove any residual particulate materials, the supernatants were further centrifuged and finally transferred to sample vials for LC-TQ/MS analysis. An equal volume of the individual samples was used to prepare the pooled plasma sample, which was used to facilitate quality control (QC), to guarantee high-quality data that were collected in batches by the high-resolution mass spectrometer and determine the reproducibility of the LC-TQ/MS system.

All trial cells cultured in 100-mm dishes were washed twice with DPBS, 1.2 mL of 80% ice-cold aqueous methanol was added, and the cells were harvested by scraping on ice. The mixed solution was transferred to a 2 mL screw-cap plastic microvial with approximately 1 g of glass beads (0.5 mm i.d.). The Mini-BeadBeater-16 (Biospec Products) is a high-energy cell disrupter, and a three-fold 2 min run (with a cooling procedure performed at the interval of each run) can disrupt cells completely. The extract was transferred to a new microcentrifuge tube and then centrifuged at 20,000 g for 10 min at 4 °C. Then, the supernatants were mixed with 800 µL of ice-cold acetonitrile and centrifuged again for protein removal. The supernatants were dried with N_2_. Dried extracts were reconstituted in 100 µL of water and then centrifuged at 20,000 g for 10 min at 4 °C. Finally, the supernatants were transferred to the LC-TQ/MS system for the metabolome assay.

Tissue samples were weighed, and 1.2 mL of 80% aqueous methanol (ice cold) was added. Then, the same protocol as that of the cells was followed to prepare analytical samples for the LC-TQ/MS-based metabolome assay; however, 1.0 mm glass beads were used for tissue disruption.

### LC-TQ/MS data acquisition

In this study, we adopted a newly developed precision-targeted metabolomics method^14^ to analyse the metabolomes of interest from trial samples (plasma, tissues, and cells). Briefly, the metabolomics method was developed with a UPLC system (1290 Infinity series, Agilent Technologies) coupled to a triple quadrupole mass spectrometer (Agilent 6495 Series Triple Quad LC/MS System). An ACQUITY UPLC HSS T3 column (2.1 mm i.d.×100 mm, 1.8 μm; Waters) was used to analyse the metabolomes of interest (**Table S1)**. The samples were placed at 10 °C with a 5 μL injection volume.

### Data analysis and visualization

First, the collected LC-TQ/MS raw data files were processed with quantitative analysis software (Agilent MassHunter Workstation System) for automatic peak recognition of the metabolomes. To guarantee the high quality of metabolome data, the peak signals of metabolites with low abundance (<10^3^) were manually excluded during the data outlook. Next, the data file from all the metabolomes was converted into a CSV file with Microsoft Excel software and uploaded onto the open-access software Metaboanalyst 4.0 (http://www.metaboanalyst.ca/MetaboAnalyst/) for data analysis and visualization, including supervised partial least-squares discriminant analysis (PLS-DA), volcano plots and unsupervised heatmap combined hierarchical cluster analysis (HCA). MS data were normalized with the total sum of all detected ions, centred and scaled using Pareto scaling, which divided each variable by the square root of the standard deviation to suppress noise interference. An unpaired t-test was used for the comparisons between the PC and NC groups to identify significantly differential metabolites (p<0.05). For biomarker discovery and performance evaluation, a receiver operating characteristic (ROC) curve was used to characterize the estimates of sensitivity (true positive rates) against 1-specificity (false-positive rates) and was performed with GraphPad Prism 8 software (GraphPad Software Inc.). ROC curves of multivariate analysis and binary logistic regression were performed using SPSS software version 21 (SPSS, Inc.). Correlation analysis of the metabolites and clinical parameters was also analysed by SPSS. All results are expressed as the mean ± SEM for replicates.

## Results

### Clinical characteristics of patients with pancreatic cancer and healthy cohort volunteers

The clinical characteristics of the study cohort are provided in **Table 1**. Fifty-eight patients who underwent resection of pancreatic cancer were staged for pancreatic cancer based on the 8th edition of the *AJCC Cancer Staging Manual*. Two patients refused further medical treatment and surgery and did not have their tumour grade recorded. In this study, the presence of metastasis was also taken into consideration for discovering potential metabolic biomarkers to diagnose the disease at an early stage and predict the high potential for metastasis.

**Table 1.** Clinical characteristics of patients with pancreatic cancer and healthy cohort volunteers

|  | PC | NC |
| --- | --- | --- |
| N | 60 | 60 |
| Age (years) | 37-79 | 38-80 |
| Gender (male/female) | 38/22 | 31/29 |
| Disease stage |  |  |
| I/II/III/IV | 21/13/13/11 |  |
| Primary/Metastatic | 36/22 |  |
| CA19-9 >37 U/ml | 50 |  |
| CEA >5 ng/ml | 16 |  |
| CA125 >35 U/ml | 17 |  |
| CA242 >20 U/ml | 38 |  |

### Plasma metabolomes identify significant metabolic alternations between patients with PC and the cohort healthy controls

Using our newly developed precision-targeted metabolomics method to comparatively analyse the plasma samples collected from the patients with pancreatic cancer (n=60) and the cohort healthy controls (n=60), substantial metabolic alternations were captured using a supervised PLS-DA model to sensitively distinguish the patients from the healthy controls. This model achieved a 73.8% rate of discrimination for Q2 and an 81.9% rate of discrimination for R2, and the score plot depicted obvious differences between the two groups, PC1 (39.2%) and PC2 (6.5%) (**Figure 2A)**. The QC samples clustered tightly together, which further confirmed the analytical reliability of the LC-TQ/MS method used to collect the metabolome data (**Figure S1**). The heatmap shows the top-differential metabolites that were observed to metabolically differentiate between the PC patients and the healthy controls (Figure 2B).

**Figure 1.**
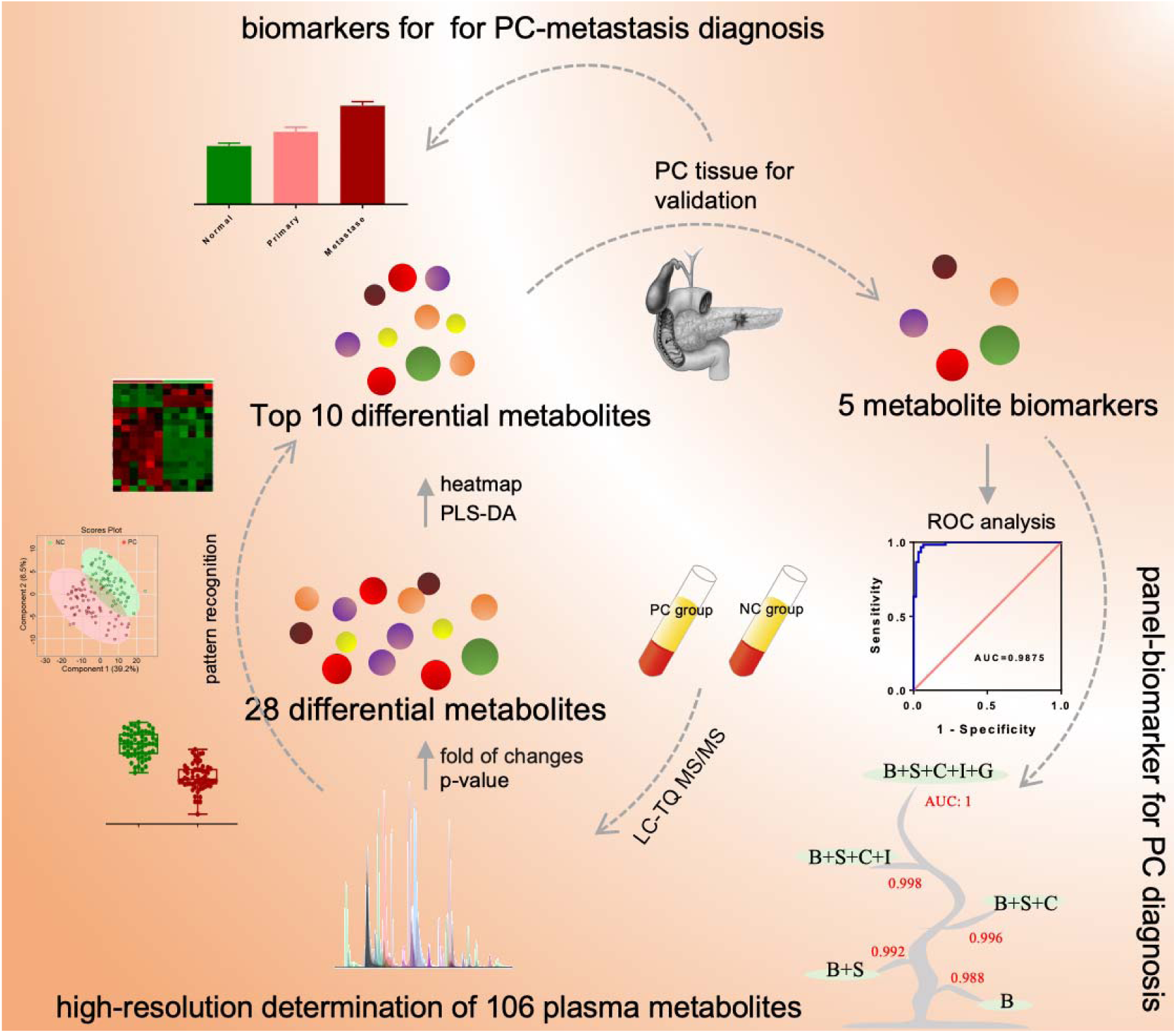
Workflow of the precision-targeted metabolomics method used to identify and validate biomarkers to diagnose the onsite progression and metastasis of pancreatic cancer in a clinical setting.

**Figure 2.**
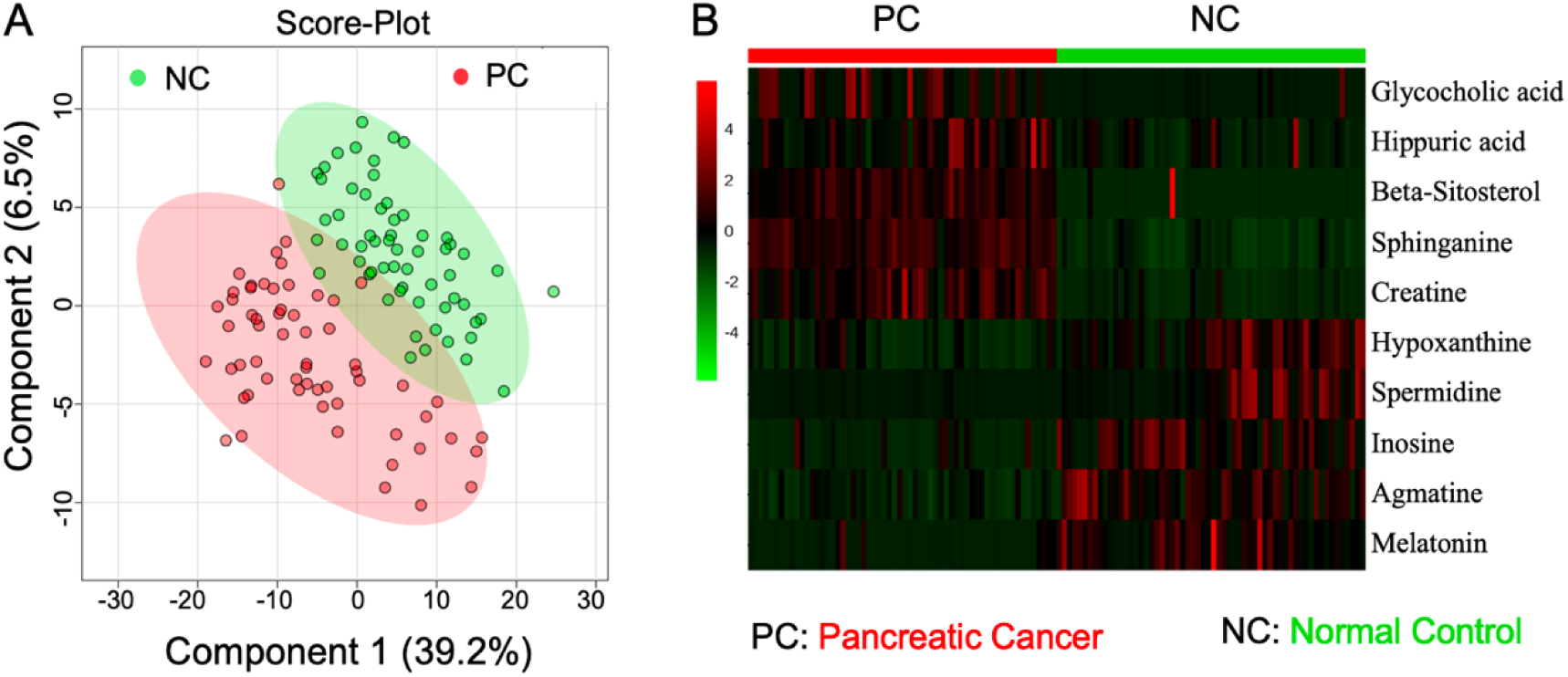
The targeted plasma metabolome assay revealed substantial metabolic differentiation of patients with PC from the cohort normal controls. (A) Score plot resulting from PLS-DA analysis of targeted metabolomes in both patients with PC and normal controls. (B) Heatmap overview of the top 10 differential metabolites whose level changes can distinguish patients with PC from normal controls.

### Differential metabolites are statistically characterized to mostly account for metabolic alternations between patients with pancreatic cancer and cohort healthy controls

The expression levels of 28 metabolites were observed to be significantly different between patients with pancreatic cancer and cohort healthy controls (FDR adjusted p-value□<□0.05; log2-fold of change>1 or <-1). The details of the differential metabolites are shown in **Table S2**. Furthermore, 18 differential metabolites were upregulated in the patients with pancreatic cancer compared to the cohort healthy controls. In addition, these differential metabolites were subjected to unsupervised heatmaps combined with hierarchical clustering analysis to obtain a pattern overview of metabolic alterations with the development of pancreatic cancer in a clinical setting. The top 10 ranked differential metabolites were precisely aligned as glycocholic acid, agmatine, melatonin, beta-sitosterol, sphinganine, hypoxanthine, spermidine, hippuric acid, creatine and inosine **(Figure 2B)**. These differential metabolites together compose the unique metabolic characteristics of pancreatic cancer, involving dysregulated metabolism in purine metabolism, glycine and serine metabolism, arginine and proline metabolism, steroid biosynthesis, sphingolipid metabolism and bile metabolism.

### Tissue metabolome assay precisely verifies high selectivity and specificity plasma biomarkers to diagnose pancreatic cancer

Because plasma metabolites are broadly biosynthesized and circulated by the whole body, only pancreatic tissue metabolites circulate into the blood, which have the capacity to assume the role of biomarkers to sensitively and specifically diagnose the physiological and pathological state of the pancreas. To further precisely verify the plasma biomarker to enable the diagnosis of pancreatic cancer, six paired samples with pancreatic cancer tissues (PCTs) and adjacent noncancerous tissues (ANTs) were collected for the specific determination of the 10 differential metabolites characterized in the last section using our targeted metabolomics method, and the expression patterns of creatine, inosine, beta-sitosterol, sphinganine and glycocholic acid agreed with those in the plasma samples (**Figure 3**). Thus, we can conclude that these differential metabolites can be regarded as biomarkers that have the capacity to diagnose PC in a clinical setting, and some have the potential to assist in the mechanistic annotation of the disease.

**Figure 3.**
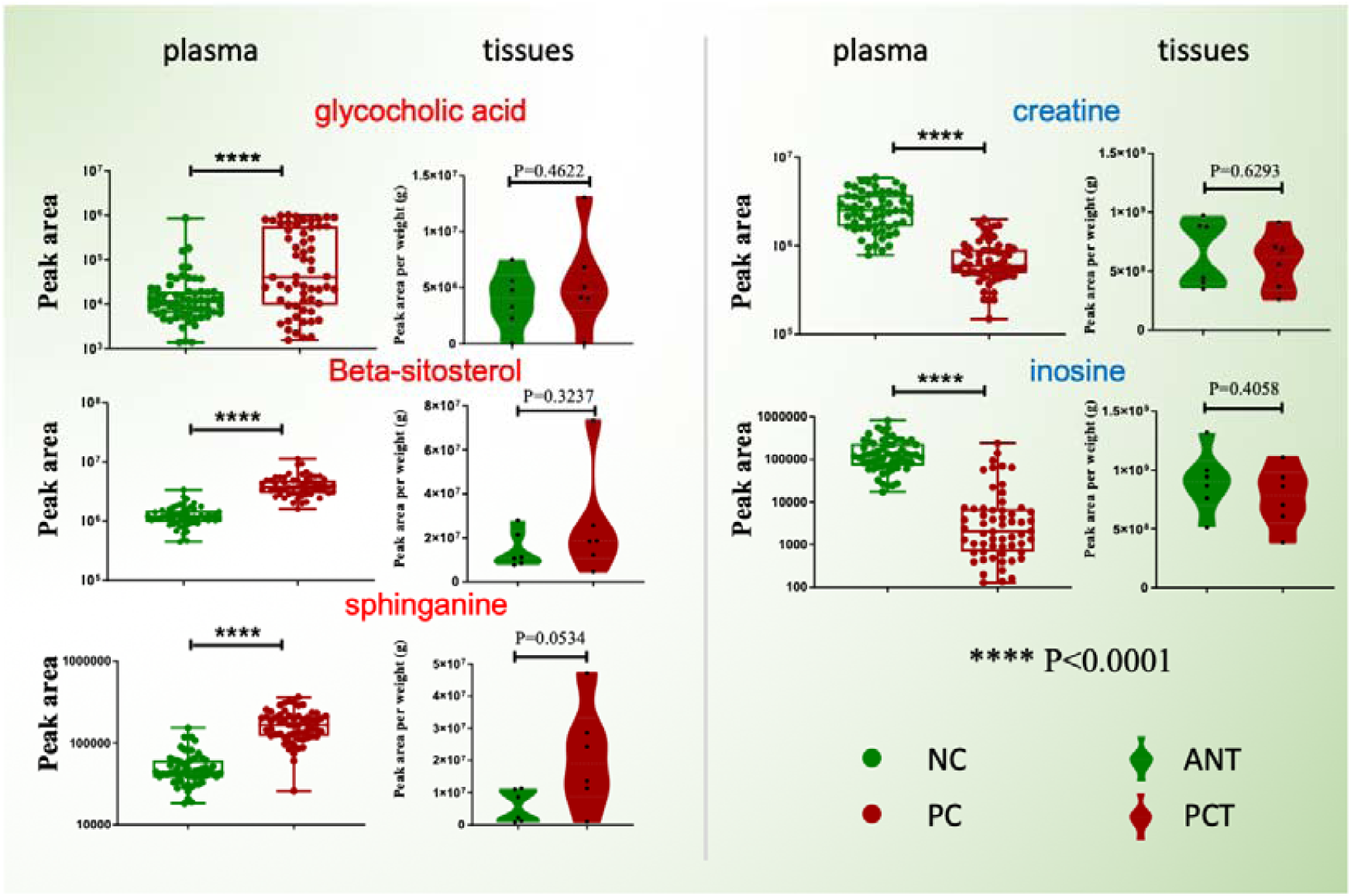
Five potential metabolite biomarkers have the capacity to diagnose pancreatic cancer in a clinical setting, and they have been verified to have agreeable changing patterns in both the tissue and plasma samples of patients with PC. ****, P<0.0001; PCT, pancreatic cancer tissue; ANT, adjacent noncancerous tissue.

### Correlation analysis between metabolite biomarkers and clinical characteristics

To investigate whether metabolic changes in the patients with pancreatic cancer were also triggered by the physiological differences among the individuals, such as sex and age in addition to disease progression, we performed Spearman correlation coefficient analysis to reveal that glycocholic acid, beta-sitosterol, sphinganine and inosine were irrelevant to gender and age, suggesting that they can be used as biomarkers to precisely diagnose patients with pancreatic cancer by neglecting the interference of age and gender. However, creatine was observed to be slightly influenced by age (*r*_s_, - 0.300) with a significant difference (*p*-value, 0.020). In addition, considering that the patients are in different stages of PC and whether nodal or distant metastases also induced metabolic differences, we found that the level of inosine in plasma is certainly correlated with the patients at the stage of nodal metastasis; this suggests that inosine is a valuable biomarker to diagnose the metastasis of PC.

### Individual differential metabolites are combined as panel biomarkers to precisely diagnose pancreatic cancer in a clinical setting

Identifying metabolite biomarkers for early disease diagnosis has a decisive impact on patient survival. To validate the selectivity and specificity of these potential biomarkers, ROC curves of individual metabolites were visualized with GraphPad Prism software. Multivariate ROC curves based on the panel metabolites, including glycocholic acid, beta-sitosterol, sphinganine, inosine and creatine, were depicted using SPSS 16.0 software. The AUCs of the five plasma metabolites were calculated to evaluate their diagnostic performance as individual biomarkers of PC. The results indicate that only glycocholic acid has a lower AUC value of 0.7064, while the other values were more than 0.95 (**Figure 4**), suggesting that creatine, inosine, beta-sitosterol and sphinganine have the capacity to be metabolite biomarkers for the efficient diagnosis of PC. To enhance the specificity and selectivity of biomarkers, we proposed a panel biomarkers strategy by combining four metabolite biomarkers as a cluster of biomarkers. Our data reveal that five biomarkers in a defined panel achieve superior diagnostic capability (AUC=1) with extremely high selectivity and specificity compared with any individual biomarker **(Figure 4).**

**Figure 4.**
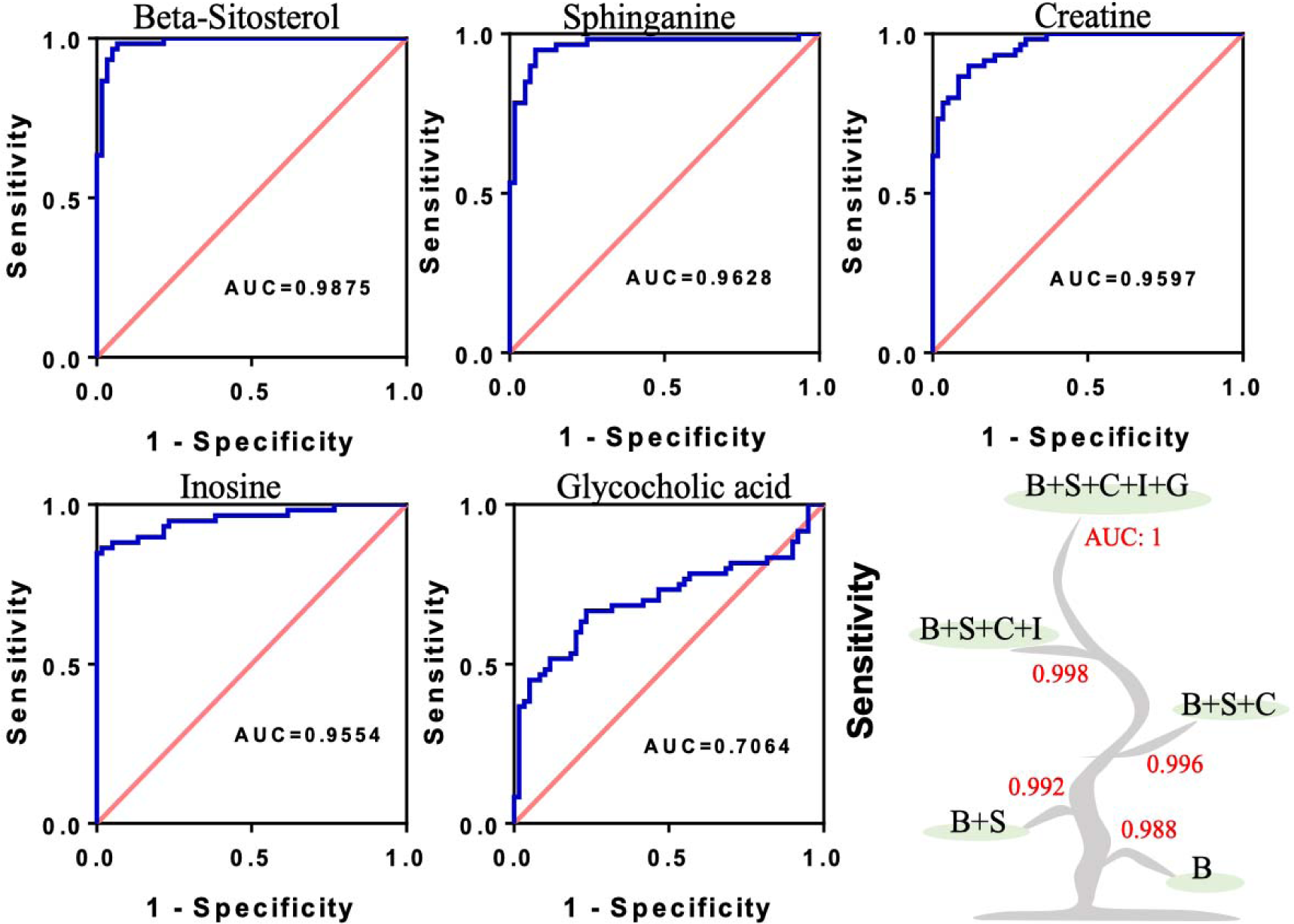
Characterization of ROC curves of univariate analysis of beta-sitosterol (B), sphinganine (S), creatine (C), inosine (I) and glycocholic acid (G), and multivariate analysis of five individual metabolites in the plasma samples collected from the PC group and NC group. AUC: Area under curve.

### The accuracy and specificity of this panel of biomarkers have been verified to be much greater than those of conventional diagnostic biomarkers for PC

CA19-9, CA125, CA242 and CEA are currently generally used as biomarkers to diagnose pancreatic cancer in the clinic. **Table 2** shows that there were no statistically significant differences in age, gender or disease stage among the PC patients. To diagnose PC, clinicians analyse the serum levels of CA125, CA19-9, CA242 and CEA to determine if the levels are higher than the cut off values of 35, 37, 20 U/ml and 5 ng/ml, respectively. However, these biomarkers lack sensitivity and specificity in the diagnosis of PC, and some PC patients cannot be properly diagnosed by measuring individual biomarkers (**Figure 5A)**. Among them, the diagnostic performance of CA19-9 is better than that of the others; almost all serum CA19-9 levels in PC patients were above the cut-off value of 37 U/ml, but 9 patients were not correctly diagnosed. To assess the diagnostic performance of our panel of biomarkers, we set these 9 special cases as the unknown group and investigated the variations in the levels of glycocholic acid, beta-sitosterol, sphinganine, inosine and creatine. PLS-DA analysis was implemented again to demonstrate that these 9 PC samples are completely discriminated from the NC cluster, which are completely clustered into the determined PC group (**Figure 5B**). In short, our panel of biomarkers were better than conventional diagnostic biomarkers by providing a highly sensitive and selective diagnostic capacity to patients with pancreatic cancer, and our panel has the potential to be used to diagnose PC.

**Table 2.**
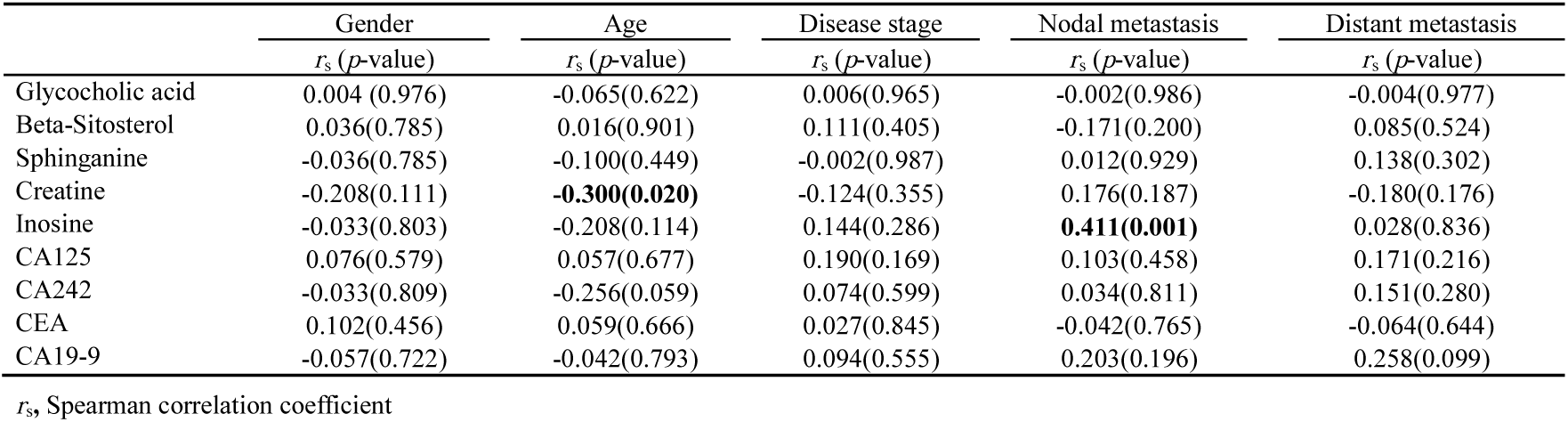
The statistics of correlation coefficients among metabolite biomarkers, clinical characteristics and conventional biomarkers of PC

**Figure 5.**
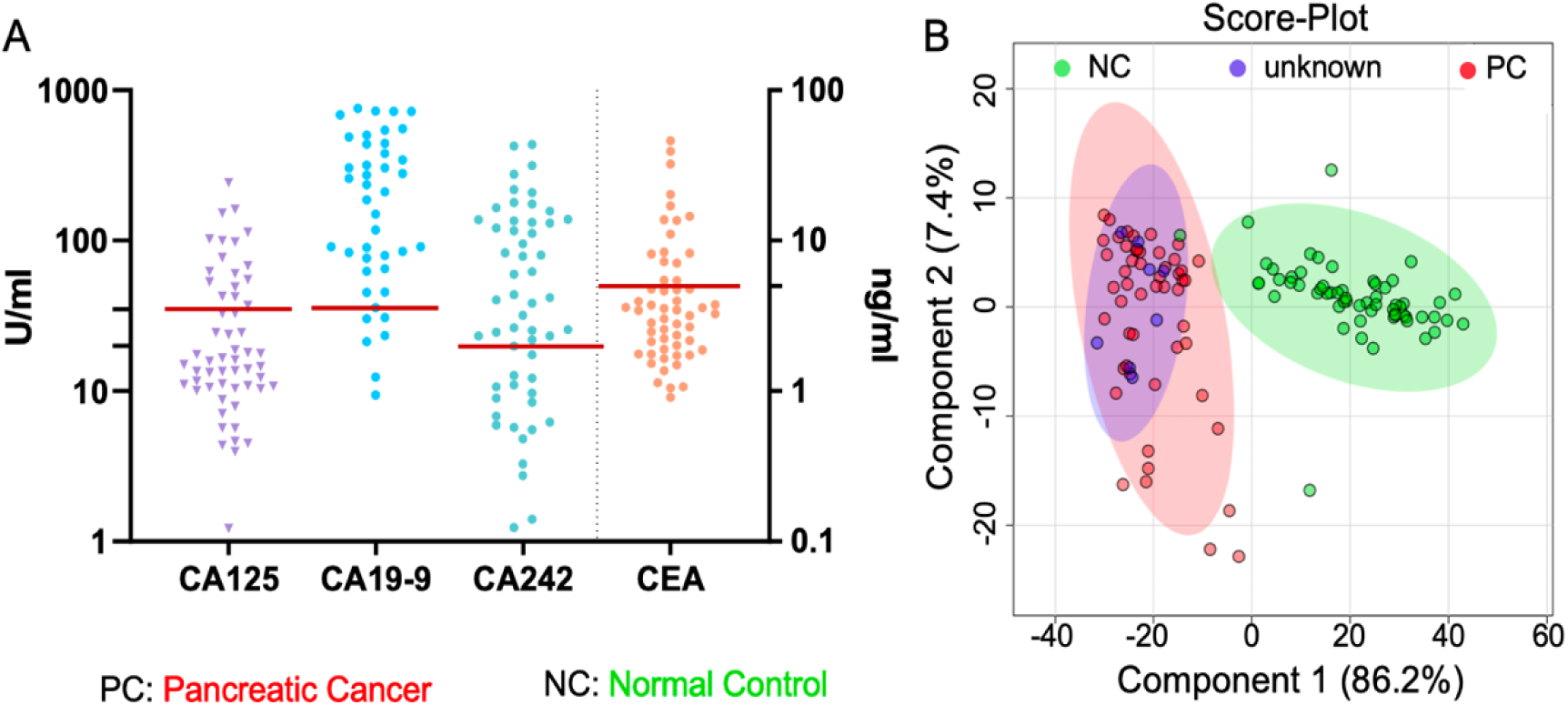
The diagnostic performance of panel biomarkers is notably higher than that of the gold standard with conventional biomarkers for PC. (A) The expression levels of serum CA125, CA19-9, CA242 and CEA in patients with PC (the cut off values are 35, 37, and 20 U/ml for CA125, CA19-9, and CA242, respectively, and 5 ng/ml for CEA). (B) The score plot resulting from PLS-DA analysis based on the expression levels of glycocholic acid, beta-sitosterol, sphinganine, inosine and creatine in the unknown group, PC group and NC group.

### Metabolite biomarkers are recognized to sensitively diagnose the progression and metastasis of PC

Lacking the proper diagnosis of cancer metastases often leads to high mortality of PC. To seek metabolite biomarkers to efficiently diagnose the progression and metastasis of PC, we employed the same metabolomics method to analyse plasma metabolomes in different stages of PC during metastases. The results demonstrated that it was difficult to identify significantly altered metabolites to distinguish PC cases in progressive stage I, stage II, stage III, and stage IV disease (**Figure 6A**), although we found slight changes with the identified biomarkers (succinic acid and gluconic acid). However, when the selected PC cases were classified into the nonmetastatic PC group and metastatic PC group, these two biomarkers were observed to efficiently diagnose PC metastases as they linearly increased compared to the nonmetastatic PC group and NC group (**Figure 6B**). Furthermore, we assessed the performance of two serum biomarkers using the selected cell lines hTERT-HPNE (normal pancreatic cells), BxPC-3 (primary PC cells), and SU.86.86 (PC metastatic cells) to determine the changes in their levels. The agreeable change patterns of the two biomarkers were visualized in both cell lines and plasma samples (**Figures 6B and 6C**). In short, succinic acid and gluconic acid should be dysregulated in PC cells and then excreted into blood; thus, plasma succinic acid and gluconic acid have the biomarker potential to sensitively diagnose PC metastases. In addition, since these two metabolite biomarkers have been observed to clearly reflect PC progression within cell lines, we confirmed the tumourigenesis of PC by deciphering their biosynthesis, utilization, metabolism and molecular regulation accordingly with cell models.

**Figure 6.**
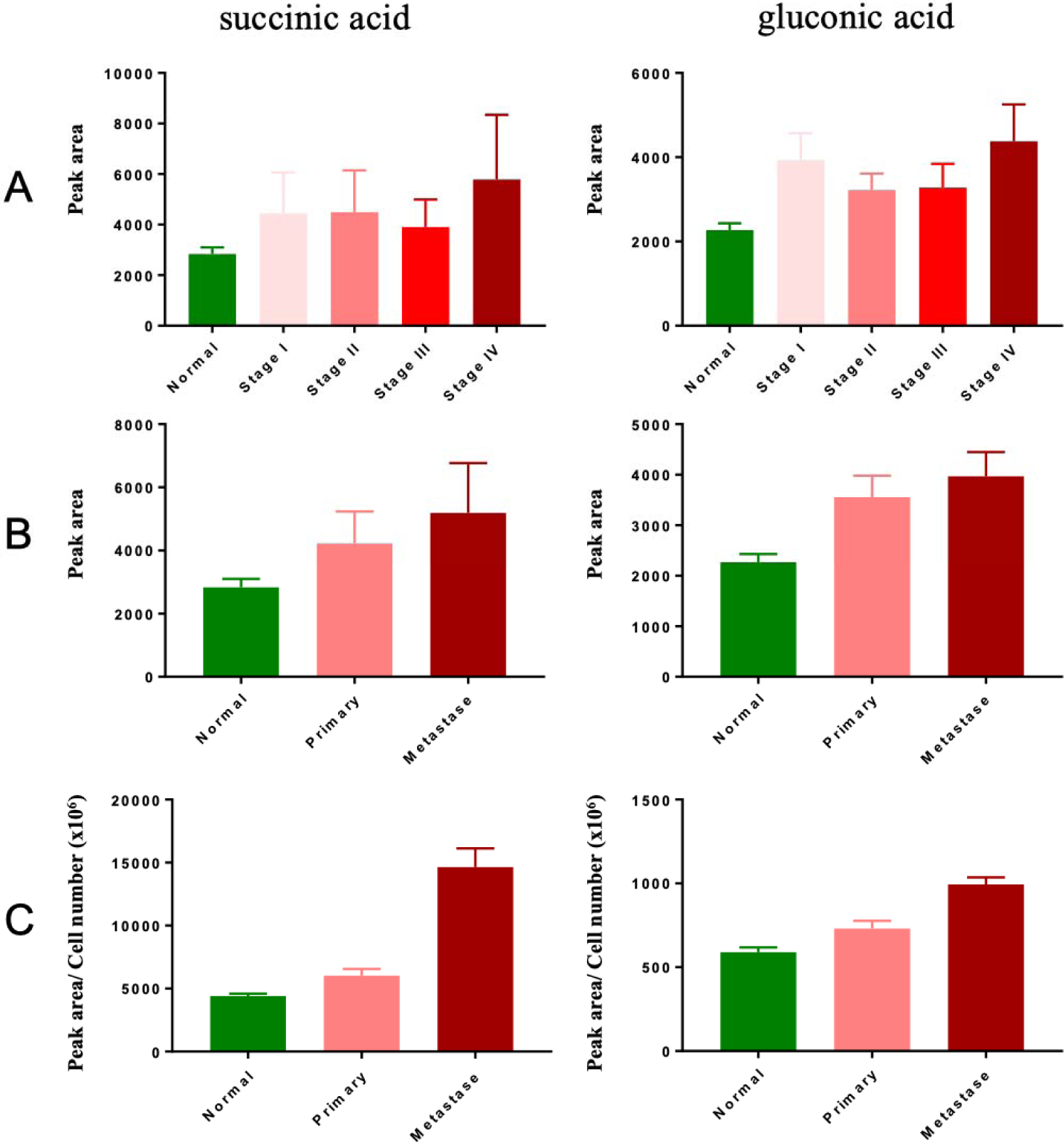
(A) Succinic acid and gluconic acid were identified as new biomarkers to monitor the progression and metastasis of pancreatic cancer in a clinical setting. Although these two biomarkers can ideally diagnose the development stage, their plasma levels are sensitively changed with the different stages. (B and C) Excitingly, they can efficiently diagnose PC in the metastatic stage and/or nonmetastatic stage, and their level changes agree in both plasma and cells, suggesting that the two biomarkers might be closely associated with the metastatic mechanisms of PC.

## Discussion

In recent years, although unexpected advances have been achieved in the treatment of cancers, the survival of PC patients remains dismal due to the severe lack of efficient diagnosis and therapeutics. Thus, the identification of molecular biomarkers is urgently needed to enhance the clinical diagnosis and monitor the metastasis of PC, which allow clinicians a new window to improve therapeutic outcomes by eventually increasing the 5-year survival rate of PC patients. Considering that cancer progression always incurs substantial metabolic costs by significantly altering the expression levels of small molecule metabolites underlying the dysregulated metabolic pathways, those altered metabolites can be defined as biomarkers to diagnose the onsite progression and metastases of PC. To verify this hypothesis, we employed the newly developed precision-targeted metabolomics method facilitated by reference compounds to analyse plasma metabolomes of interest in both patients with pancreatic cancer and healthy controls. This method is unlike the untargeted metabolomics method, which has a great challenge in valuable data discovery and precision identification of metabolite biomarkers ^18^.

Through a metabolome assay of plasma samples collected from PC patients and the cohort healthy controls, the identities of creatine, inosine, beta-sitosterol, sphinganine, and glycocholic acid were confirmed, and their level changes can significantly distinguish PC patients from healthy controls. Additionally, their specificity for PC was verified by their determination in paired PC tissues and adjacent normal pancreatic tissues, by which we confirmed that they should be produced by pancreatic cells and then excreted into the blood, as they share agreeable change patterns in both plasma and tissue. This result suggests that these differential metabolites have the potential to be diagnostic biomarkers for PC. To evaluate biomarker performance, ROC curve analysis was performed to reveal that creatine, inosine, beta-sitosterol, and sphinganine have much higher sensitivity and specificity than the known glycocholic acid to diagnose pancreatic cancer. Glycocholic acid is a secondary bile acid, and particularly high levels of bile acids have been implicated in the pathogenesis of liver diseases^19^. One study suggested that glycocholic acid and taurochenodeoxycholic acid can be used as phenotypic biomarkers in cholangiocarcinoma^20^. In this study, glycocholic acid in plasma was observed to be positively correlated with pancreatic cancer, suggesting that bile acids might also be closely associated with pancreatic cancer. Beta-sitosterol and sphinganine in plasma were upregulated compared with healthy controls. Beta-sitosterol is a main dietary phytosterol found in plants that may induce apoptosis in human breast cancer cells and can be used for cancer prevention and therapy^21,22^. However, the plasma beta-sitosterol concentration in PC patients closely depends on oral dietary intake, and the upregulation in PC patients may be attributed to lower utilization. Sphinganine is an intermediate of sphingoid base biosynthesis^23,24^. Upregulation of sphinganine suggests that sphingolipid metabolism is hampered in PC progression. This affects the downstream metabolites ceramide and sphingosine-1-phosphate, which have played important roles in regulating the death and survival of cancer cells ^25,26^. In addition, sphingosine 1-phosphate has been indicated to confer an important function in regulating glucose-stimulated insulin secretion in pancreatic beta cells^27^.

The expression levels of creatine and inosine in plasma were significantly decreased in PC patients. A decrease in inosine in PC plasma has been reported^28,29^, but the biological significance of inosine in the tumourigenesis of PC is still poorly understood^30^. Creatine was also found to significantly decrease with age. Due to the lower muscle mass of elderly people, creatine biosynthesis may be a particular burden compared with younger individuals^31^. Disturbances in creatine metabolism are generally associated with diseases of skeletal muscle, the heart, the brain, and the kidneys^32^. Rare research has reported the correlation of creatine biosynthesis with pancreatic cancer. Therefore, our data reveal that the strong correlation between creatine and PC could open a new window for improving diagnosis and therapeutics. Although our promising result was to identify individual biomarkers that can efficiently diagnose PC, their diagnostic performance has not reached an optimal level, especially for the diagnostic specificity of PC. To improve this issue, we defined a panel biomarker strategy by accumulating different individual biomarkers into one combinational panel. Excitingly, a panel biomarker strategy enables improved diagnostic accuracy and specificity of pancreatic cancer when we combined five biomarkers into one panel to facilitate the diagnosis of PC.

Then, we further explored biomarkers using the same metabolomics method to efficiently diagnose the progression and metastases of PC in a clinical setting. The results confirmed that succinate and gluconate, which are important intermediates of the TCA cycle and pentose phosphate pathway (PPP), have the capability to monitor the progression and metastases of PC. Radioactive gluconate has been used in positron emission tomography (PET) analysis as an indicator of cancerous lesions for proper localization of the lesions and accurate assessment of malignancy^33^. This must be evidence to support the clinical feasibility of gluconate to diagnose the progression and metastasis of PC. Succinate is a key modulator of the hypoxic response. According to the Warburg effect^34^, most cancer cells employ aerobic glycolysis rather than oxidative phosphorylation to synthesize energy ^35^. Proliferating cells divert glucose metabolites into anabolic processes, while glutamine is provided as a carbon source for the TCA cycle^36^. Succinate, as an important intermediate of the TCA cycle, is also intertwined with metabolites of the glutamine metabolic pathway and plays a crucial role in tumourigenesis^37^. The PPP is a major glucose catabolic pathway and a recent study revealed that the PPP pathway can generate cellular NADPH to fuel tumour cells for rapid proliferation^38^. It is unclear how cancer cells coordinate fuel biosynthesis to support the rapid growth of tumours. Importantly, key functional metabolism will be implemented in the near future, with emphasis on the mechanistic exploration of PC tumourigenesis by deciphering the metabolic regulation of biomarker-associated pathways (**Figure 7**).

**Figure 7.**
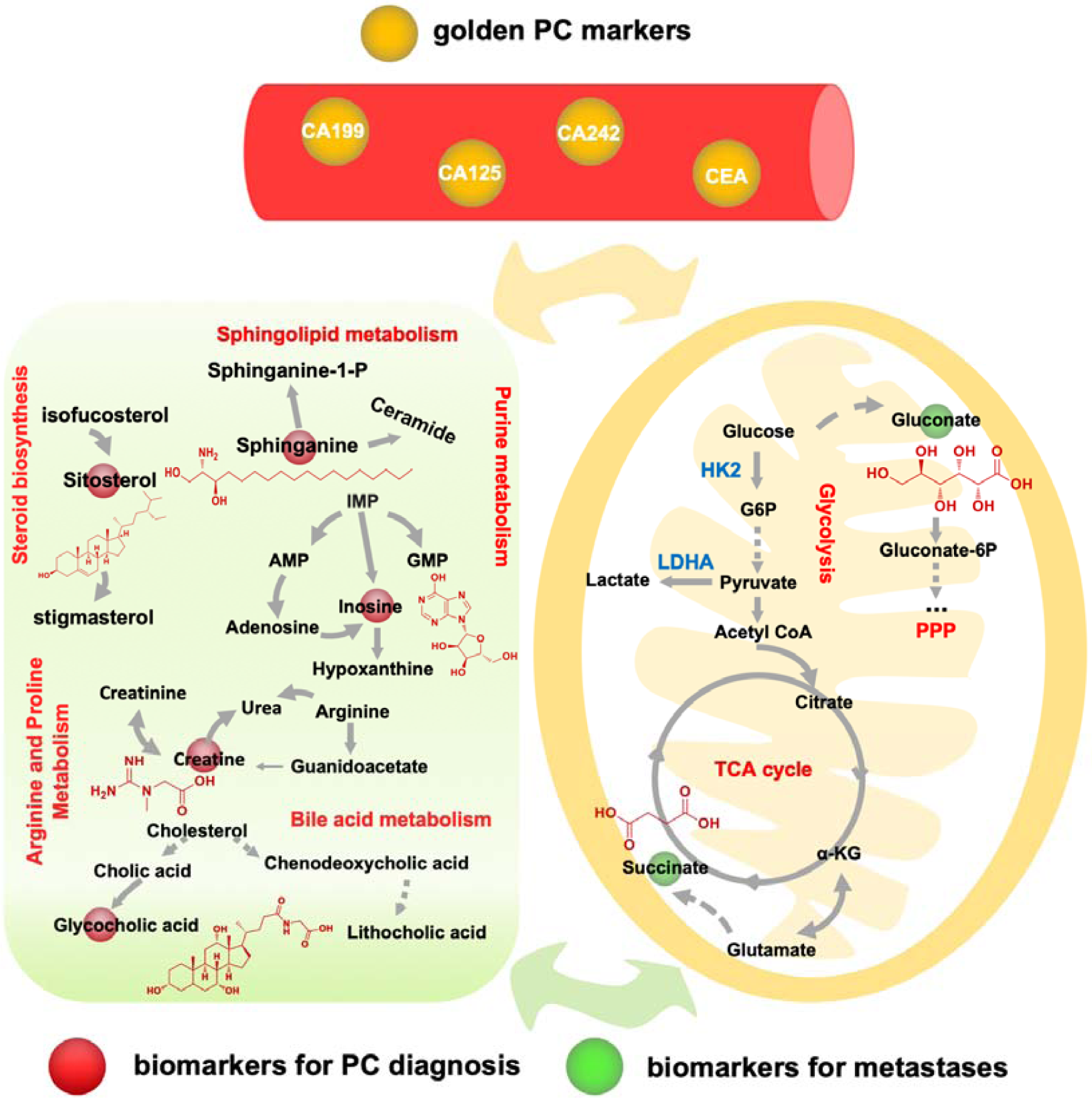
The most affected metabolic pathways are annotated during the onsite, progression and metastasis of pancreatic cancers and the pentose phosphate pathway, TCA cycle, amino acid metabolism and lipid metabolism are dysregulated, which are supposed to be implicated in the tumorigenesis of PC.

## Conclusion

Using our newly developed precision-targeted metabolomics method, we successfully identified plasma biomarkers (creatine, inosine, beta-sitosterol, sphinganine and glycocholic acid) that can precisely diagnose pancreatic cancer by efficiently distinguishing patients with PC from their cohort of healthy controls, which have been verified with the differential metabolome assay of tissues and cell lines associated with PC. We proposed a panel biomarker strategy to achieve excellent diagnostic performance with much higher sensitivity and selectivity than each individual biomarker, and excitingly, the panel biomarker was verified to demonstrate extremely high accuracy compared to the conventional PC biomarkers (CA19-9, CA125, CA242 and CEA), which have lower specificity and selectivity. In addition, we also identified two plasma biomarkers (succinic acid and gluconic acid) that can efficiently diagnose the progression and metastasis of PC, which also underlie the tumourigenesis of PC. In short, regarding the pathogenic latency that often leads to the late diagnosis and high mortality of PC, our innovations in new biomarker discovery to efficiently diagnose the onsite progression and metastases of PC by which we can precisely diagnose PC at an early stage pertains to the capacity to monitor PC metastases from a metabolic perspective and enables clinicians to achieve PC prevention and treatment in different developmental stages. Eventually, the optimized therapeutic window shall be left to the clinician to largely improve therapeutic outcome and enhance the 5-year survival rate in the patient community.

## Acknowledgments

This work was supported by the National Key R&D Program of China (No. 2017YFC1308600 and 2017YFC1308605), the National Natural Science Foundation of China Grants (No. 81274175 and 31670031), the Startup Funding for Specialized Professorship Provided by Shanghai Jiao Tong University (No. WF220441502).

## Figure legends

**Table S1.** The RPLC gradient elution program

| Time (min) | A % | B % |
| --- | --- | --- |
| 0 | 98 | 2 |
| 2 | 98 | 2 |
| 10 | 65 | 35 |
| 12 | 20 | 80 |
| 14 | 2 | 98 |

A: H<sub>2</sub>O with 0.1% formic acid; B: ACN with 0.1% formic acid

**Table S2.** The volcano plot analysis identified 28 differential metabolites

|  | Metabolites | log2 (FC) | p adjusted |
| --- | --- | --- | --- |
| Downregulated | Spermidine | -4.0651 | 1.89E-06 |
|  | Inosine | -2.8267 | 2.28E-10 |
|  | Hippuric acid | -2.6358 | 8.02E-07 |
|  | D-Fructose | -2.2355 | 4.48E-11 |
|  | Glucose | -2.2106 | 4.48E-11 |
|  | Hypoxanthine | -1.4677 | 2.96E-10 |
|  | Creatine | -1.2744 | 2.41E-13 |
|  | L-Aspartic acid | -1.1912 | 1.30E-06 |
| Upregulated | Taurocholic acid | 6.4908 | 0.00030081 |
|  | Taurochenodeoxycholic acid | 4.6698 | 0.00057625 |
|  | Taurodeoxycholic acid | 4.6698 | 0.00057625 |
|  | Tauroursodeoxycholic acid | 4.2093 | 0.00015019 |
|  | Glycocholic acid | 3.5659 | 6.87E-06 |
|  | Melatonin | 3.3842 | 5.01E-16 |
|  | Taurohyodeoxycholic acid | 2.5742 | 0.0024263 |
|  | Malonic acid | 2.4232 | 7.79E-05 |
|  | Sphinganine | 2.3758 | 1.84E-14 |
|  | Beta-Sitosterol | 2.3436 | 3.51E-17 |
|  | Glycochenodeoxycholic acid | 1.8914 | 0.00027029 |
|  | Uridine 5'-monophosphate | 1.8904 | 3.80E-08 |
|  | Caproic acid | 1.574 | 7.07E-08 |
|  | Gluconic acid | 1.3496 | 2.50E-09 |
|  | Succinic acid | 1.2766 | 0.0020688 |
|  | Dehydrocholic acid | 1.0692 | 1.02E-14 |
|  | Methylmalonic acid | 1.0492 | 0.0030922 |
|  | Agmatine | 1.0205 | 1.41E-05 |

**Figure S1.**
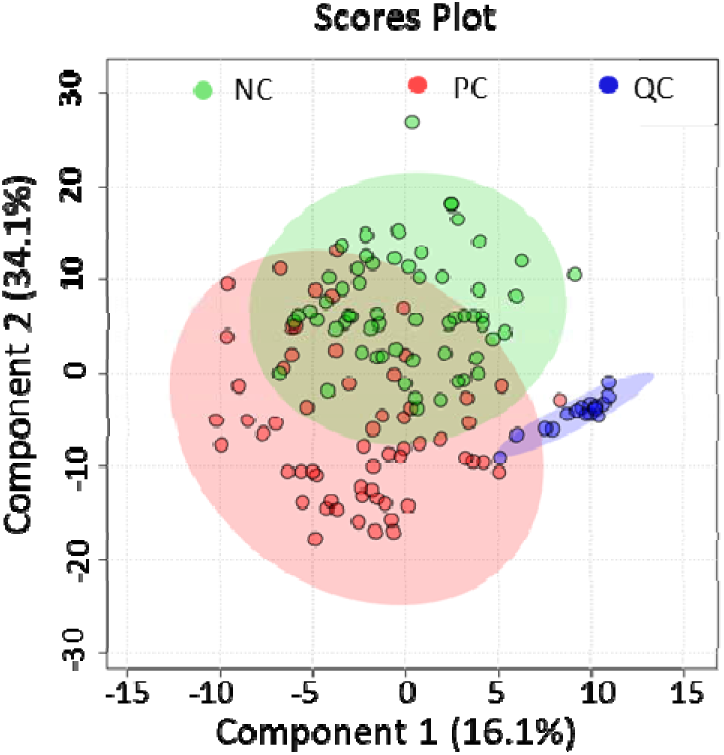
The partial least-squares discriminant analysis (PLS-DA) result score plot shows that QC samples are clustered tightly into one pattern during the batch running of metabolomics samples, suggesting that the LC-TQ/MS system has acceptable stability and reproducibility, and that the collected metabolome data are highly reliable for biochemical annotation.

## References

1 Wen, C. L. et al. An allosteric PGAM1 inhibitor effectively suppresses pancreatic ductal adenocarcinoma. Proceedings of the National Academy of Sciences of the United States of America, doi: 10.1073/pnas.1914557116 (2019).

2 Winter, J. M. et al. Survival after resection of pancreatic adenocarcinoma: results from a single institution over three decades. Annals of surgical oncology 19, 169–175, doi: 10.1245/s10434-011-1900-3 (2012).

3 Pothula, S. P. et al. Key role of pancreatic stellate cells in pancreatic cancer. Cancer letters 381, 194–200, doi: 10.1016/j.canlet.2015.10.035 (2016).

4 Luo, G. et al. Potential Biomarkers in Lewis Negative Patients With Pancreatic Cancer. Annals of surgery 265, 800–805, doi: 10.1097/sla.0000000000001741 (2017).

5 Berger, A. W. et al. A Blood-Based Multi Marker Assay Supports the Differential Diagnosis of Early-Stage Pancreatic Cancer. Theranostics 9, 1280–1287, doi: 10.7150/thno.29247 (2019).

6 Abe, T. et al. Gene Variants That Affect Levels of Circulating Tumor Markers Increase Identification of Patients with Pancreatic Cancer. Clinical gastroenterology and hepatology : the official clinical practice journal of the American Gastroenterological Association, doi: 10.1016/j.cgh.2019.10.036 (2019).

7 Saiki, Y. & Horii, A. Molecular pathology of pancreatic cancer. Pathol Int 64, 10–19, doi: 10.1111/pin.12114 (2014).

8 Bailey, P. et al. Genomic analyses identify molecular subtypes of pancreatic cancer. Nature 531, 47–52, doi: 10.1038/nature16965 (2016).

9 Roe, J. S. et al. Enhancer Reprogramming Promotes Pancreatic Cancer Metastasis. Cell 170, 875–888 e820, doi: 10.1016/j.cell.2017.07.007 (2017).

10 Johnson, C. H. et al. Metabolomics guided pathway analysis reveals link between cancer metastasis, cholesterol sulfate, and phospholipids. Cancer Metab 5, 9, doi: 10.1186/s40170-017-0171-2 (2017).

11 Gebregiworgis, T. et al. Glucose Limitation Alters Glutamine Metabolism in MUC1-Overexpressing Pancreatic Cancer Cells. Journal of proteome research 16, 3536–3546, doi: 10.1021/acs.jproteome.7b00246 (2017).

12 Son, J. et al. Glutamine supports pancreatic cancer growth through a KRAS-regulated metabolic pathway. Nature 496, 101-+, doi: 10.1038/nature12040 (2013).

13 Nishiumi, S. et al. Serum metabolomics as a novel diagnostic approach for pancreatic cancer. Metabolomics 6, 518–528, doi: 10.1007/s11306-010-0224-9 (2010).

14 Luo, X., Zhang, A., Wang, X. & Lu, H. UHPLC/MS based large-scale targeted metabolomics method for multiple-biological matrix assay. bioRxiv, 642496, doi: 10.1101/642496 (2019).

15 Hui, S. et al. Glucose feeds the TCA cycle via circulating lactate. Nature 551, 115–118, doi: 10.1038/nature24057 (2017).

16 Chan, A. K., Bruce, J. I. & Siriwardena, A. K. Glucose metabolic phenotype of pancreatic cancer. World journal of gastroenterology 22, 3471–3485, doi: 10.3748/wjg.v22.i12.3471 (2016).

17 Yuan, C. et al. Circulating Metabolites and Survival Among Patients With Pancreatic Cancer. Journal of the National Cancer Institute 108, djv409, doi: 10.1093/jnci/djv409 (2016).

18 Cui, L., Lu, H. & Lee, Y. H. Challenges and emergent solutions for LC-MS/MS based untargeted metabolomics in diseases. Mass Spectrom Rev 37, 772–792, doi: 10.1002/mas.21562 (2018).

19 Loftfield, E. et al. Prospective investigation of serum metabolites, coffee drinking, liver cancer incidence, and liver disease mortality. Journal of the National Cancer Institute, doi: 10.1093/jnci/djz122 (2019).

20 Song, W. S. et al. Discovery of glycocholic acid and taurochenodeoxycholic acid as phenotypic biomarkers in cholangiocarcinoma. Sci Rep 8, 11088, doi: 10.1038/s41598-018-29445-z (2018).

21 Awad, A. B., Roy, R. & Fink, C. S. Beta-sitosterol, a plant sterol, induces apoptosis and activates key caspases in MDA-MB-231 human breast cancer cells. Oncology reports 10, 497–500.

22 Muti, P. et al. A plant food-based diet modifies the serum beta-sitosterol concentration in hyperandrogenic postmenopausal women. The Journal of nutrition 133, 4252–4255, doi: 10.1093/jn/133.12.4252 (2003).

23 Knapp, M., Baranowski, M., Lisowska, A. & Musiał, W. Decreased free sphingoid base concentration in the plasma of patients with chronic systolic heart failure. Advances in Medical Sciences 57, 100–105, doi: https://doi.org/10.2478/v10039-011-0057-4 (2012).

24 Fan, Y. et al. Comprehensive Metabolomic Characterization of Coronary Artery Diseases. J Am Coll Cardiol 68, 1281–1293, doi: 10.1016/j.jacc.2016.06.044 (2016).

25 Cartier, A. & Hla, T. Sphingosine 1-phosphate: Lipid signaling in pathology and therapy. Science (New York, N.Y.) 366, doi: 10.1126/science.aar5551 (2019).

26 Ogretmen, B. Sphingolipid metabolism in cancer signalling and therapy. Nature reviews. Cancer 18, 33–50, doi: 10.1038/nrc.2017.96 (2018).

27 Cantrell Stanford, J. et al. Sphingosine 1-phosphate (S1P) regulates glucose-stimulated insulin secretion in pancreatic beta cells. J Biol Chem 287, 13457–13464, doi: 10.1074/jbc.M111.268185 (2012).

28 Kang, C. M. et al. Postoperative serum metabolites of patients on a low carbohydrate ketogenic diet after pancreatectomy for pancreatobiliary cancer: a nontargeted metabolomics pilot study. Scientific reports 9, 16820–16820, doi: 10.1038/s41598-019-53287-y (2019).

29 Urayama, S., Zou, W., Brooks, K. & Tolstikov, V. Comprehensive mass spectrometry based metabolic profiling of blood plasma reveals potent discriminatory classifiers of pancreatic cancer. Rapid Communications in Mass Spectrometry 24, 613–620, doi: 10.1002/rcm.4420 (2010).

30 Alseth, I., Dalhus, B. & Bjørås, M. Inosine in DNA and RNA. Current Opinion in Genetics & Development 26, 116–123, doi: https://doi.org/10.1016/j.gde.2014.07.008 (2014).

31 Brosnan, J. T., da Silva, R. P. & Brosnan, M. E. The metabolic burden of creatine synthesis. Amino Acids 40, 1325–1331, doi: 10.1007/s00726-011-0853-y (2011).

32 Wyss, M. & Kaddurah-Daouk, R. Creatine and creatinine metabolism. Physiological reviews 80, 1107–1213, doi: 10.1152/physrev.2000.80.3.1107 (2000).

33 Mycielska, M. E. et al. Potential Use of Gluconate in Cancer Therapy. Front Oncol 9, 522–522, doi: 10.3389/fonc.2019.00522 (2019).

34 Liberti, M. V. & Locasale, J. W. The Warburg Effect: How Does it Benefit Cancer Cells? Trends Biochem Sci 41, 211–218, doi: 10.1016/j.tibs.2015.12.001 (2016).

35 Bhattacharya, B., Mohd Omar, M. F. & Soong, R. The Warburg effect and drug resistance. Br J Pharmacol 173, 970–979, doi: 10.1111/bph.13422 (2016).

36 Ying, H. et al. Oncogenic Kras Maintains Pancreatic Tumors through Regulation of Anabolic Glucose Metabolism. Cell 149, 656–670, doi: https://doi.org/10.1016/j.cell.2012.01.058 (2012).

37 Jiang, S. & Yan, W. Succinate in the cancer-immune cycle. Cancer letters 390, 45–47, doi: 10.1016/j.canlet.2017.01.019 (2017).

38 Yao, P. et al. Evidence for a direct cross-talk between malic enzyme and the pentose phosphate pathway via structural interactions. J Biol Chem 292, 17113–17120, doi: 10.1074/jbc.M117.810309 (2017).

